# Cox-nnet: an artificial neural network method for prognosis prediction on high-throughput omics data

**DOI:** 10.1101/093021

**Authors:** Travers Ching, Xun Zhu, Lana X. Garmire

## Abstract

Artificial neural networks (ANN) are computing architectures with massively parallel interconnections of simple neurons and has been applied to biomedical fields such as imaging analysis and diagnosis. We have developed a new ANN framework called Cox-nnet to predict patient prognosis from high throughput transcriptomics data. In over 10 TCGA RNA-Seq data sets, Cox-nnet achieves a statistically significant increase in predictive accuracy, compared to the other three methods including Cox-proportional hazards (Cox-PH), Random Forests Survival and CoxBoost. Cox-nnet also reveals richer biological information, from both pathway and gene levels. The outputs from the hidden layer node can provide a new approach for survival-sensitive dimension reduction. In summary, we have developed a new method for more accurate and efficient prognosis prediction on high throughput data, with functional biological insights. The source code is freely available at github.com/lanagarmire/cox-nnet.

## Introduction

Artificial Neural networks (ANNs) were developed in 1943 in order to model the activity of neurons ^1^. In recent years, ANNs have caught renewed attention, thanks to the increased parallel computing power and the promise of deep learning ^2^. The original ANN extension of Cox Regression was not designed to handle high throughput input data^3^. Some recent attempts using ANNs to high dimensional survival data simplified the regression problem as either a binary classification or fitting discrete variables of survival time through binning, leading to loss of accuracy ^4,5^. Another study used time as an additional input in order to predict patient survival or censoring status^6^, with the potential to overfit if the survival time and censoring time are correlated. To avoid all these issues, we herein leverage the neural network extension of Cox regression by a high-performance and easy-to-use package, particularly fit for high dimensional data.

Besides Cox-nnet, some other modeling methods exist to predict patient survival. The standard method is Cox proportional hazards (Cox-PH) regression, a semi-parametric and generalized linear model with an exponential link function^7^. Another method, CoxBoost ^8^, is an iterative “boosting” method modified from the Cox-PH model. In each boosting iteration, it refits the parameters by maximizing the penalized likelihood function. Rather than using L2 ridge regression common in Cox-PH, the number of boosting iterations is used as the complexity parameter in CoxBoost and optimized via cross-validation (CV) ^8^. Random Forests Survival (RF-S) is another ensemble, non-linear method ^9^. It combines many bootstrapped decision trees in order to reduce the variance in the model, and then calculates a weighted average of all the decision trees. Unlike Cox-PH and CoxBoost, RF-S does not use the log likelihood function to determine the fitness of the model. Instead, it predicts estimated survival times, and uses Harrel's C-Index, a score that measures the correct ranking of individuals ^9^.

The new software package we have developed here, named Cox-nnet, advances the ANN extension of Cox regression for survival prediction on high-throughput data. The caliber of this package is manifested in several aspects. First, it has improved technical performance in terms of both accuracy and speed. In 4 comparison with the other methods mentioned above (Cox-PH, RF-S and CoxBoost), Cox-nnet has better overall predictive accuracy. It is also optimized on graphics processing unit (GPU) with at least an order of computational speed-up over the central processing unit (CPU), making it a compelling new tool to predict disease prognosis in the era of precision medicine. Second, Cox-nnet utilizes feature importance scores based on the partial derivatives of gene features selected by the model, so that the relative importance of the genes to prognosis outcome can be directly assessed. Thirdly, the hidden layer node structure in ANN can be harnessed to reveal much richer information of featuring genes and biological pathways, compared to the Cox-PH method. Overall, Cox-nnet is a desirable survival analysis method with both excellent predictive power and usage to gain biological functions related to prognosis.

## Methods

### The Cox model

The Cox-PH model is a log-linear model that estimates individual hazard, i.e., an instantaneous measure of the likelihood of an event, based on a set of features. The hazard is given by the equation:

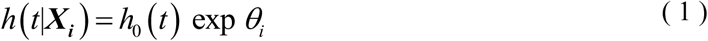

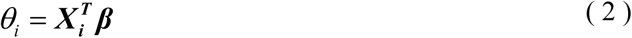

Where *θ_i_* is the log hazard ratio for patient *i*. The partial likelihood is represented by the following formula:

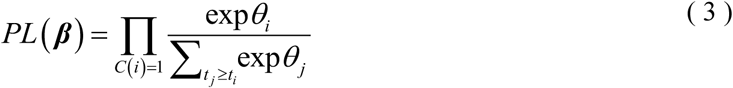

Where *C* (*i*) is the censoring status of a patient, and *C are found by minimizing the cost function* (*i*) = 0 if the patient was censored or 1 if the patient died or had a recurrence event, etc. The partial log-likelihood is used as the cost function:

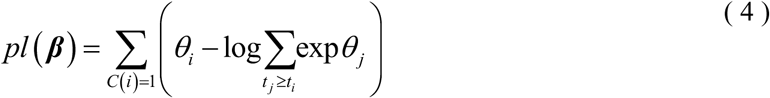

In a Cox model with L2 ridge regression, a penalty term is added which is proportional to the L2 norm of the coefficients. The cost function is minimized to find the best coefficients for the model:

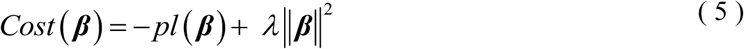

where the tuning parameter *λ* is determined by maximizing CV.

The cross-validated performance metric may be Harrel's concordance index (C-index) ^10^ or the “cross-validated partial likelihood” ^11^. Since the contribution of each patient in the partial likelihood is determined only in the context of all the other patients, the cross-validated partial likelihood is calculated subtracting full partial likelihood from the training set in the CV. In the k-th iteration of a K-fold CV, the optimal coefficients ***β***̂_*λ,k*_ are found by minimizing the cost function on the training sub-samples. If *pl_k_* (***β***̂_*λ,k*_) is the partial likelihood of the training sub-samples, and *pl* (***β***̂_*λ,k*_)is the partial likelihood of the full dataset, then the cross-validated partial likelihood is the sum of differences:

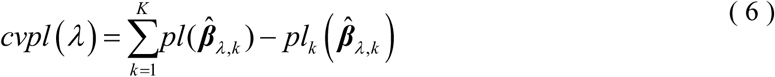

### ANN extension of Cox regression

The ANN extension of Cox regression (Cox-nnet) is a neural network whose output layer is replaced by a Cox model. In a Cox-nnet model with one input layer of *J* input features and one hidden layer composed of *H* hidden nodes, the linear predictor is replaced by the outputs of the hidden layer:

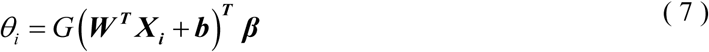

Where ***W*** is the coefficient weight matrix between the input and hidden layer with the size H x J, *b* is the bias term for each hidden node and *G* is the activation function (applied element-wise on a vector). Subsequently, the ridge regression cost function is modified to:

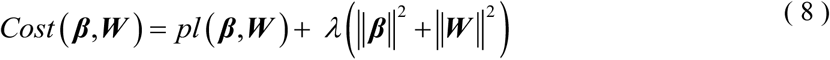

In this manuscript, the tanh activation function is used, as it results in faster training time compared to the sigmoid activation ^12^. The tanh function is:

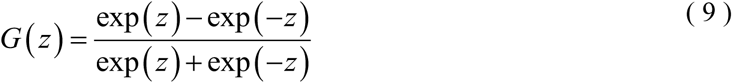

In addition to ridge regularization, we also employ dropout regularization^13^. In this approach, nodes are removed during each training iteration with probability 1-p. During evaluation, output from the nodes are multiplied by p. The optimal dropout parameter, p, is determined through cross-validation on the training set. Dropout regularization has been shown to reduce overfitting and improve performance over other regularization schemes^13^.

The source code of cox-nnet can be found at: github.com/lgarmire/cox-nnet, and can be installed through the Python Package Index (PyPI). Documentation of package can be found at lgarmire.github.io/cox-nnet/docs.

### Implementation in Python with Theano

We implement Cox-nnet using a feed forward, back propagation network with gradient descent. The partial log likelihood is usually written as a double conditional sum (equation 4). To avoid the computational inefficiency of calculating the partial log likelihood (equation 4) using two nested for loops, we convert it into a formulation of matrix operations and basic sums. First we define an indicator matrix ***R*** with elements:

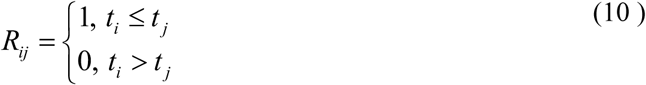

We also define an indicator vector ***C*** with elements given by the censoring of each patient. An operation using ***R*** replaces the conditional sum over *t_j_* ≥ *t_i_*, and an operation using ***C*** replaces the conditional sum over *C* (*i*) = 1 in equation 4. In Theano, the partial log likelihood is:

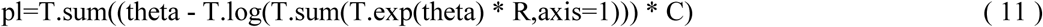

### Model evaluation

To evaluate the performance of all methods in comparison, we trained each model on 80% of the samples for each dataset (chosen randomly) and evaluated the performance on the 20% holdout test set. The output of Cox-PH, Cox-nnet and CoxBoost are the log hazard ratios (i.e., Prognosis Index, or PI) for each patient. The hazard ratio describes the relative risk of a patient compared to a non-parametric baseline. On the other hand, the output of RF-S is an estimation of the survival time for each patient.

We use C-index and log-ranked p-value based on dichotomization of the hold-out test data of the holdout test data to measure the performance of each model. The C-index is a measure of how well the model prediction corresponds to the ranking of the survival data ^14^. It is calculated for censored survival data, which evaluates a value between 0 and 1, with 0.5 equivalent to a random process. The C-index can be computed as a summation over all events in the dataset, whereby patients with a higher survival time and lower log hazard ratios (and conversely patients with a lower survival time but higher log hazard ratios) are considered concordant. The C-index is a measure of concordance of the data with the model prediction. To calculate the log-ranked p-value, a PI cutoff threshold is used to dichotomize the patients in the data set into higher and lower risk groups, similar to our earlier report ^15,16^. A log-ranked p-value is then computed to differentiate the Kaplan-Meier survival curves between the higher vs. lower risk groups. In this report, we used the median log hazard ratio as the cutoff threshold.

### Feature evaluation

For computing the importance of a feature in Cox-nnet, we use a method of partial derivatives (PaD) ^17,18^. For each patient, we compute the partial derivatives of each input with respect to the linear output of the model (e.g., the log hazard ratio). The average of the partial derivatives for each input across all patient samples is calculated as the feature score.

### Datasets

In order to evaluate the performance of Cox-nnet, we analyzed 10 TCGA datasets which were combined into a pan-cancer dataset. The TCGA datasets included the following cancer types: Bladder Urothelial Carcinoma (BLCA), Breast invasive carcinoma (BRCA), Head and Neck squamous cell carcinoma (HNSC), Kidney renal clear cell carcinoma (KIRC), Brain Lower Grade Glioma (LGG), Liver hepatocellular carcinoma (LIHC), Lung adenocarcinoma (LUAD), Lung squamous cell carcinoma (LUSC), Ovarian serous cystadenocarcinoma (OV) and Stomach adenocarcinoma (STAD). RNA-Seq expression and clinical data were downloaded from the Broad Institute GDAC ^19^. Overall survival time and censoring information were extracted from the clinical follow-up data. Raw count data were normalized using the DESeq2 R package ^20^ and then log-transformed. Datasets were selected from TCGA based on the following criteria: > 300 samples with both RNASeq and survival data and > 50 survival events. In total, 5031 patient samples were used (see Table S1 for a patient tabulation by individual dataset).

## Results

### Cox-nnet structure and optimization

Cox-nnet is the neural network extension of the Cox-PH model. We created a package suitable for high dimensional datasets using the Theano math library in Python. The neural network model used in this paper is shown in Figure 1 and an overview of modules in the Cox-nnet package is shown in Figure S1. As a proof of concept, the current ANN architecture is composed of three layers: one input layer, one fully connected hidden layer and an output “Cox regression” layer. The output layer of Cox-nnet replaces the linear predictors in the standard Cox-PH model. Many other functions are implemented to improve the usability of the package, including CVSearch, CVProfile, CrossValidation, and TrainCoxMlp. CVSearch, CVProfile, CrossValidation are methods that perform CV to find the optimal regularization parameter. TrainCoxMlp performs optimization of coefficients on the regularized partial likelihood function. The optimization strategies include momentum gradient descent ^21^, Nesterov accelerated gradient ^22^ and Ada Delta ^23^. A comparison of these descent methods is shown in Figure S2A, where Nesterov accelerated gradient method achieved the best efficiency based on TCGA kidney renal clear cell carcinoma (KIRC) data. Moreover, this package can be run on multiple threads or a Graphics Processing Unit (GPU), and it achieves slightly faster training time compared to Random Forest and CoxBoost (Figure S2B). Thus, Cox-nnet is a modern software implementation that can achieve efficient computational time.

**Figure 1.**
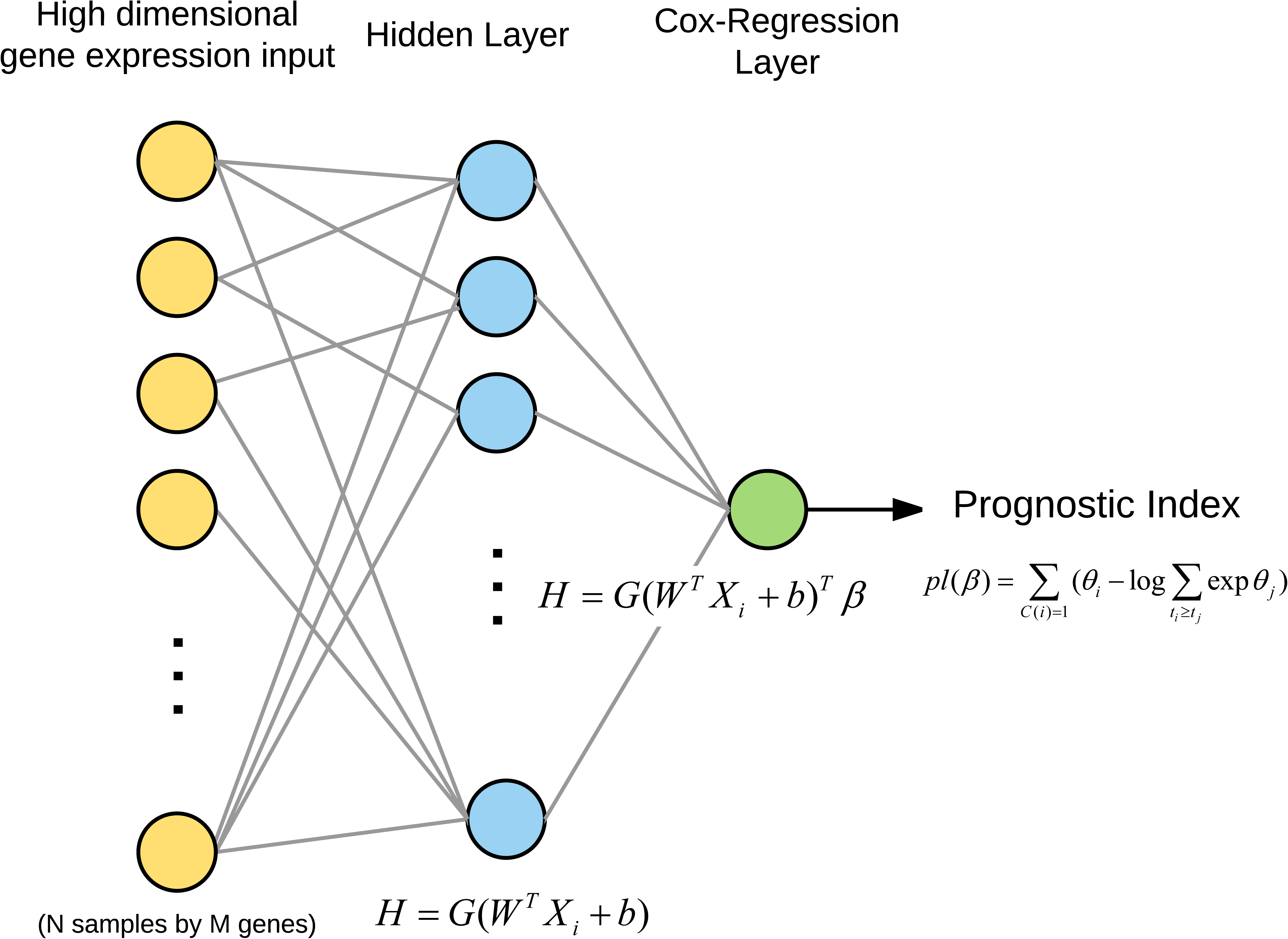
An overview of the neural network architecture used in this study.

### Performance comparison of survival prediction methods

We compared four methods, including Cox-nnet, Cox-PH, CoxBoost and RF-S, on 10 datasets from The Cancer Genome Atlas (TCGA), which were selected based on having at least 50 death events (Table S1). For each dataset, we trained the model on 80% of the randomly selected samples and determined the regularization parameter using 5-fold CV on the training set. We used two types of regularizations, L2 ridge regularization (also known as weight decay) and dropout regularization. We evaluated the performance on the remaining 20% holdout test set. Two metrics are used to evaluate the performance of the model. The first one is Harrell's concordance index (C-index) calculated for censored survival data ^10,24^. It evaluates the relative ordering of the samples and ranges between 0 and 1, with 0.5 equivalent to a random process. The second metric is the log-ranked p-value from Kaplan-Meier survival curves of two different survival risk groups. This is done by using the median threshold of Prognosis Index (PI), the output of Cox-nnet, to dichotomize the patients into higher and lower risk groups, similar to our earlier reports ^15,16,24^ A log-ranked p-value is then computed to differentiate the Kaplan-Meier survival curves from these two groups.

The comparison of C-indices among the four methods over the 10 TCGA data is shown in Figure 2A. Overall, Cox-nnet has higher predictive accuracy over the other three methods, regardless of the regularization method. Cox-PH performs the second best, followed by CoxBoost and RF-S in descending order (Figure 2B). The comparison of log-ranked p-values on the dichotomized survival risk groups is shown in Figure S3. Generally, log-ranked p-values in the 10 TCGA datasets are more significant in Cox-nnet, compared to other methods. However, the dichotomization of patients ignores the differences within each dichotomized group, thus the resulting log-ranked p-values are less consistent than C-indices on the same data.

**Figure 2.**
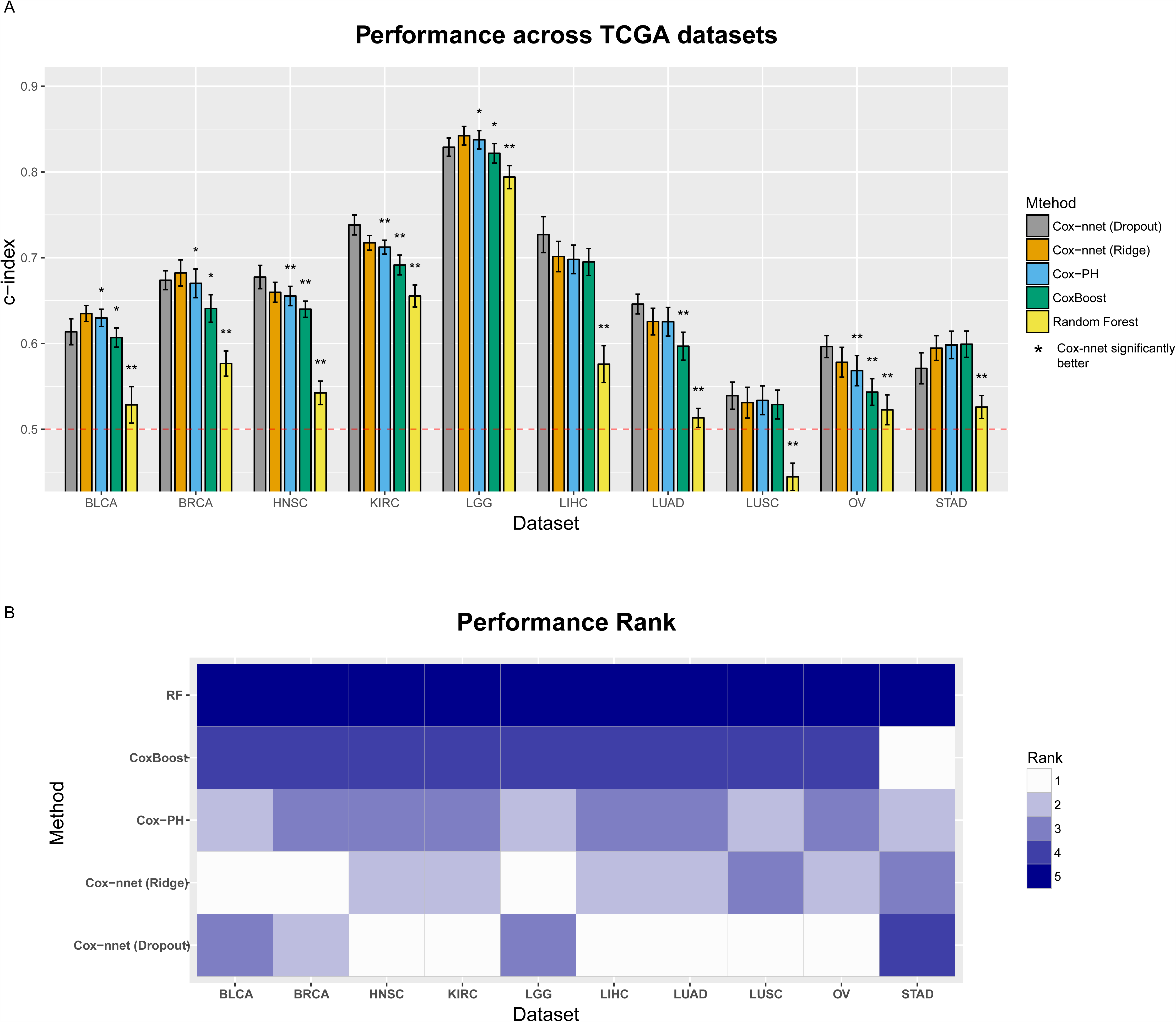
A. Barplot of the C-index of the 10 TCGA datasets using four prognosis-predicting methods (Cox-nnet, CoxBoost, Cox-PH and RF-S). Each dataset was randomly split into 80% training and 20% testing sets. B. Heatmap of the performance rank of each dataset.

### Biological relevance of hidden layer nodes of Cox-nnet

To explore the biological relevance of the hidden nodes of Cox-nnet, we used the TCGA KIRC dataset as an example. We first extracted the contribution of each hidden node to the PI score for each patient (Figure 3A). The contribution was calculated as the output value of each hidden node weighted by the corresponding coefficient at the Cox regression output layer. As expected, the value of the hidden nodes strongly correlated to the PI score. However, there is still significant heterogeneity among the nodes, suggesting that individual nodes may reflect different biological processes. We hypothesize that the top nodes may serve as surrogate features to discriminate patient survival. To explore this idea, we selected the top 20 nodes with the highest variances, and presented the patients PI scores using t-SNE, a popular method to enhance the separation among samples^25^. The nodes represent a dimension reduction of the original data and clearly discriminate samples by their PI scores (Figure 3B). In contrast, the top 20 principle components obtained from principal component analysis (PCA) in combination with t-SNE fail to separate the patient samples (Figure 3B). This drastic difference demonstrates that the nodes in Cox-nnet effectively capture the survival information, and the top node PI scores can be used as features for dimension reduction in survival analysis.

**Figure 3.**
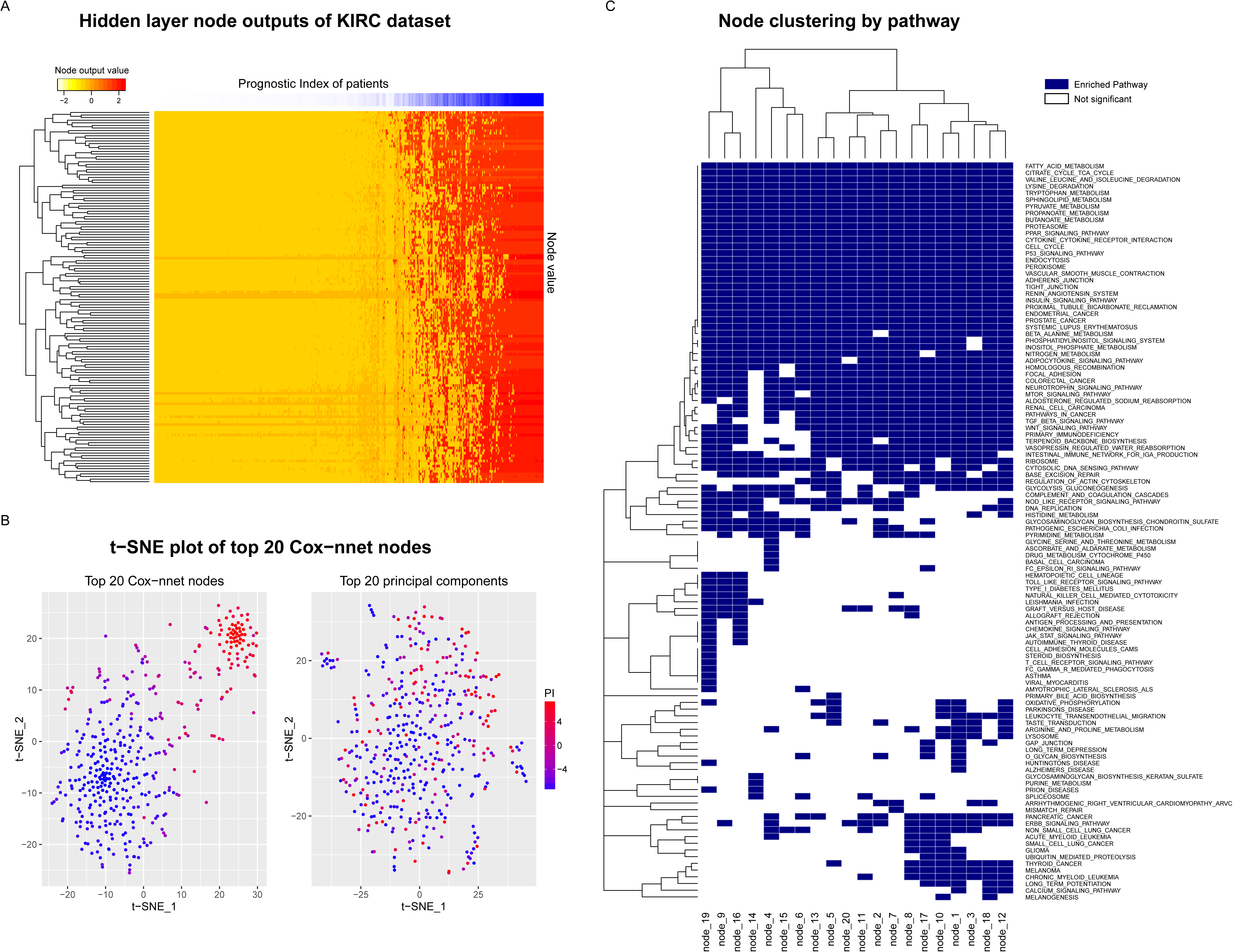
A. Hidden node output of the TCGA KIRC dataset. B. t-SNE plot of the top 20 hidden nodes and the top 20 principal components in PCA analysis. C. Gene Set Enrichment Analysis: significantly enriched KEGG pathways of the top 20 hidden nodes (adjusted p-value < 0.05).

To further explore the biological relevance of the top 20 hidden nodes, we conducted Gene Set Enrichment Analysis (GSEA)^26^ using KEGG pathways ^27^. We calculated significantly enriched pathways using gene correlation to the output score of each node (Figure 3C and Table S2), and compared these enriched pathways to those from GSEA of the Cox-PH model (Table S3). To calculate statistical significance of the pathways, we performed 10,000 permutations, followed by multiple hypothesis testing with Benjamini Hochberg adjustment. A total of 110 (out of 187) significantly enriched pathways (Table S2) were identified in at least one node, including seven pathways enriched in all 20 nodes that were not found by the Cox-PH method (Table 1). In contrast, Cox-PH only identified 30 significantly enriched pathways using the same significance threshold. Among the seven pathways, the P53 signaling pathway stands out as an important biologically relevant pathway (Figure 4 and Figure S4), since it was shown to be highly prognostic of patient survival in kidney cancer^28^.

**Figure 4.**
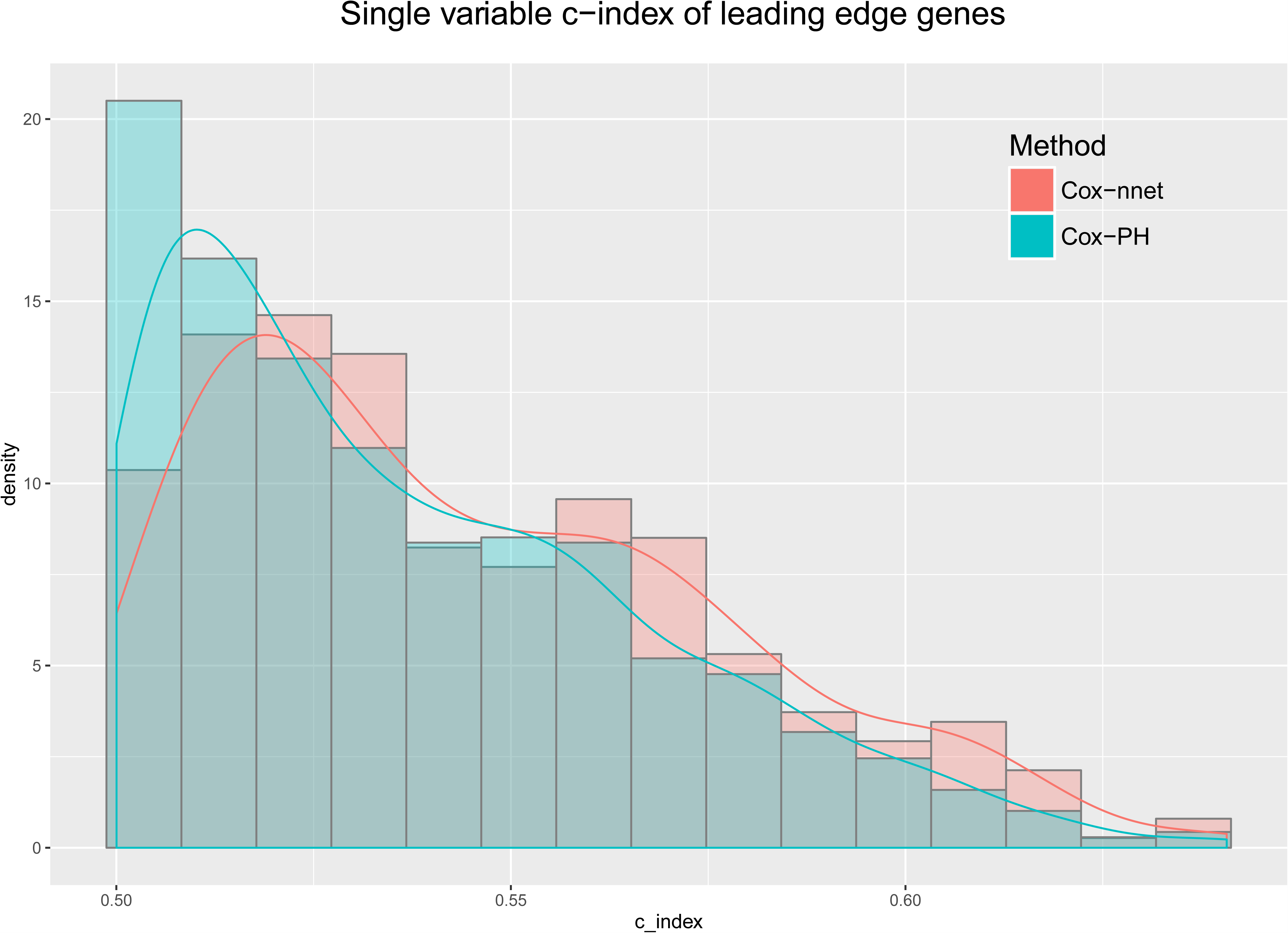
Single variable C-index scores of the leading edge genes from Cox-nnet and Cox-PH. Cox-nnet has significantly higher C-index scores (p = 5.79e-5).

**Table 1.** Cox-nnet node-associated pathways. Significantly enriched pathways from common to all 20 hidden nodes that are not found in the Cox-PH Gene Set Enrichment Analysis (Adjusted p < 0.05).

| Pathway | P.value | P.adjusted | Nodes |
| --- | --- | --- | --- |
| KEGG adherens junction | 0.000 | 0.001 | 1-20 |
| KEGG endocytosis | 0.000 | 0.001 | 1-20 |
| KEGG insulin signaling pathway | 0.000 | 0.001 | 1-20 |
| KEGG lysine degradation | 0.000 | 0.003 | 1-20 |
| KEGG p53 signaling pathway | 0.000 | 0.003 | 1-20 |
| KEGG pyruvate metabolism | 0.000 | 0.001 | 1-20 |
| KEGG sphingolipid metabolism | 0.001 | 0.005 | 1-20 |

Next, we estimated the predicative accuracies of the leading edge genes (LEG) enriched in the KEGG pathways from Cox-nnet vs. those enriched in Cox-PH model. We used the C-index of each LEG, obtained from single-variable analysis (Figure 4). Collectively, LEGs from Cox-nnet have significantly higher C-index scores (p = 5.79e-05) than those from Cox-PH, suggesting that Cox-nnet has selected more informative features. In order to visualize these gene level and pathway level differences between Cox-nnet and Cox-PH, we reconstructed a bipartite graph between LEGs for Cox-nnet or feature genes (for Cox-PH) and their corresponding enriched pathways (Figure 5). Besides P53 pathway mentioned earlier that is specific to Cox-nnet, several other pathways, such as insulin signaling pathway, endocytosis and adherens junction, also have many more genes enriched in Cox-nnet. Among them, some have been previously reported to relevant to renal carcinoma development and prognosis, such as CASP9^29^, TGFBR2^30^, KDR (VEGFR)^31^. These results demonstrate that Cox-nnet model reveals richer biological information than Cox-PH.

**Figure 5.**
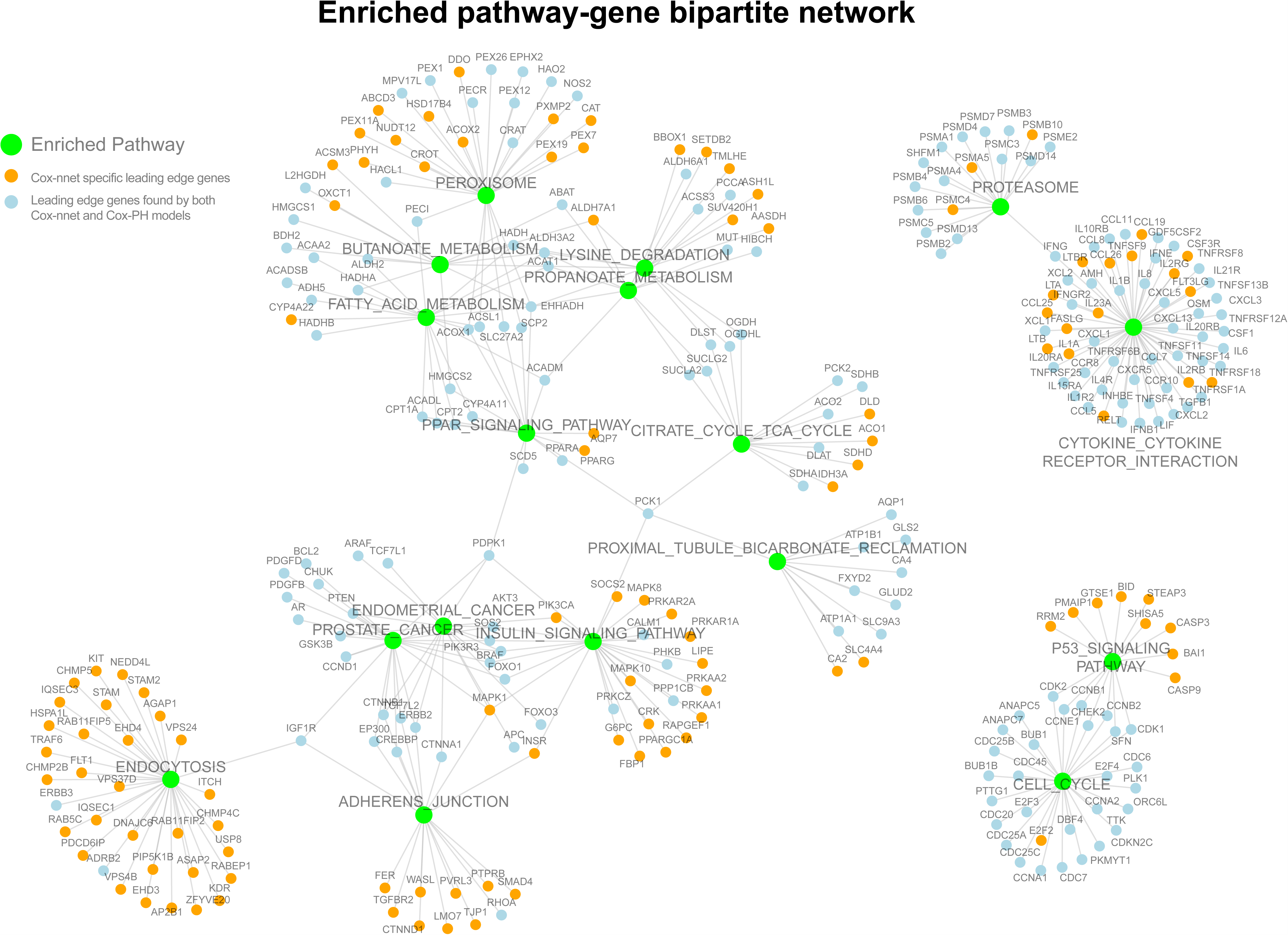
Enriched pathway-gene bipartite network from the leading edge genes and significantly enriched pathways. Significantly enriched pathways common to all 20 hidden nodes are labeled in green. Leading edge genes found uniquely in Cox-nnet are labeled in orange, and genes found in both Cox-nnet and Cox-PH are labeled in blue.

To further examine the importance of each gene relative to the survival outcome, we calculated the averaged partial derivative (PaD) of each input gene feature over all patients, with respect to the linear output of the model (e.g., the log hazard ratio). As demonstrated by the LEGs in seven common pathways of all nodes in Cox-nnet, the feature importance scores produce stronger biological insight (Figure S4). For example, the feature importance for the BAI1 gene in the P53 pathway is much higher in the Cox-nnet model compared to the Cox-PH model. Corresponding to our finding, the BAI gene family was found to be involved in several types of cancers including renal cancer^32 33 34 35^. BAI1 acts as an inhibitor to angiogenesis and is transcriptionally regulated by P53 ^36^. Its expression level was significantly decreased in tumor vs. normal kidney tissue, and was even lower in advanced stage renal carcinoma^35^. Mice kidney cancer models treated with BAI1 showed slower tumor growth and proliferation ^37^. Additionally, the MAPK1 gene (also known as ERK2) has a much higher feature importance score in Cox-nnet compared to Cox-PH, and is annotated in the Adherens Junction pathway as well as the Insulin Signalling Pathway found by Cox-nnet. MAPK1 is one of the key kinases in intra-cellular transduction, and was found constitutively activated in renal cell carcinoma ^38^. Drugs inhibiting the MAPK cascade have been targeted for development^39^.

## Discussion

In this report, we have implemented Cox-nnet, a new non-linear ANN method, to predict patient survival from high throughput omics data. Cox-nnet is an improved, modern alternative to the standard Cox-PH regression, as demonstrated by increased performance for survival prediction and the capabilities to explore more deeply the biological information.

First, through in-depth comparison of 10 TCGA RNA-Seq, Cox-nnet achieves overall statistically significant improvements over Cox-PH on its predictive accuracy, as measured by C-indices. Interestingly, the ensemble-based method RF-S consistently ranks worse than Cox-nnet and Cox-PH. Because RF-S bootstraps both samples and features for individual trees, many uninformative features in each tree may be chosen for node splitting in particularly high dimensional datasets, leading to a decrease in overall accuracy ^40^. In contrast, the dropout and L2-regularization approach used by both Cox-nnet and Cox-PH can prune out uninformative features.

Second, Cox-nnet can reveal a lot richer biological information than Cox-PH. This is manifested both at the pathway and gene levels. The hidden nodes in the Cox-nnet model have distinct expression patterns, and can serve as surrogate features for survival-sensitive dimension reduction. Many more significant KEGG pathways are enriched which correlate with top nodes in Cox-nnet, as compared to those from the Cox-PH model. A critical pathway for renal cancer development, P53 pathway, is only enriched by Cox-nnet but not Cox-PH model in TCGA KIRC. Other pathways, including insulin signaling pathway, endocytosis and adherens junction, have many more genes enriched by Cox-mmet. Moreover, leading edge genes (LEGs) obtained from these KEGG pathways enriched by Cox-nnet (which are a fraction of the gene features considered by the model) have collectively higher associations with survival.

As a promising new predictive method for prognosis, the current Cox-nnet implementation has some limitations. Its architecture includes 3-layer ANN, and it is possible to incorporate other more sophisticated architecture into the model, such as including more layers of neurons. A convolutional neural network approach using convolutional and pooling layers could also be used, as those reported in processing imaging or other types of positional data ^41^. Additionally, it is possible to embed *a priori* biological pathway information into the network architecture, e.g., by connecting genes in a pathway to a common node in the next hidden layer of neurons. In the future, we plan to further analyze how different neural network architectures affect the performance of Cox-nnet and compare the biological insights from the various models.

## Author Contributions

LXG and TC envisioned the project and designed the work. TC coded the project and conducted the analysis. XZ provided insight and discussion on neural networks. All authors have read, revised and approved the final manuscript.

## Competing financial interests

The author(s) declare no competing financial interests.

## Acknowledgements

This research was supported by grants K01ES025434 awarded by NIEHS through funds provided by the trans-NIH Big Data to Knowledge (BD2K) initiative (www.bd2k.nih.gov), P20 COBRE GM103457 awarded by NIH/NIGMS, R01 LM012373 awarded by NLM, R01 HD084633 awarded by NICHD, and Hawaii Community Foundation Medical Research Grant 14ADVC-64566 to L.X. Garmire.

---

Figure S1. An overview of the structure, methods and classes in Cox-nnet package.

Figure S2. A: comparison of descent methods on the TCGA KIRC dataset. The change in cost function is evaluated over 100,000 iterations for three methods: gradient descent, momentum gradient descent and the Nesterov accelerated gradient. B: Training time comparing CPU training time vs. GPU training time on the same dataset.

Figure S3. Log-rank p-value bar plot of the 10 TCGA datasets.

Figure S4.Variable importance of the common leading edge genes of enriched KEGG pathways.

Table S1. Tabulation of TCGA patients by individual dataset.

Table S2. Significantly enriched pathways from the Cox-PH method (p < 0.05).

Table S3. Significantly enriched pathways from the Cox-nnet method (p < 0.05).

## References

1 McCulloch, W. S. & Pitts, W. A logical calculus of the ideas immanent in nervous activity. The bulletin of mathematical biophysics 5, 115–133 (1943).

2 Jones, N. (Nature Publishing Group MACMILLAN BUILDING, 4 CRINAN ST, LONDON N1 9XW, ENGLAND, 2014).

3 Faraggi, D. & Simon, R. A neural network model for survival data. Statistics in medicine 14, 73–82 (1995).

4 Petalidis, L. P. et al. Improved grading and survival prediction of human astrocytic brain tumors by artificial neural network analysis of gene expression microarray data. Molecular cancer therapeutics 7, 1013–1024 (2008).

5 Chi, C.L., Street, W. N. & Wolberg, W. H. in AMIA Annual Symposium Proceedings. 130 (American Medical Informatics Association).

6 Joshi, R. & Reeves, C. in Proceedings of the eighteenth international conference on systems engineering. 179–184.

7 Therneau, T. M. & Grambsch, P. M. Modeling survival data: extending the Cox model. (Springer Science & Business Media, 2000).

8 Binder, H. CoxBoost: Cox models by likelihood based boosting for a single survival endpoint or competing risks. R package version 1 (2013).

9 Ishwaran, H., Kogalur, U. B., Blackstone, E. H. & Lauer, M. S. Random survival forests. The Annals of Applied Statistics, 841–860 (2008).

10 Koziol, J. A. & Jia, Z. The concordance index C and the Mann—Whitney parameter Pr (X> Y) with randomly censored data. Biometrical Journal 51, 467–474 (2009).

11 van Houwelingen, H. C., Bruinsma, T., Hart, A. A., van’t Veer, L. J. & Wessels, L. F. Cross-validated Cox regression on microarray gene expression data. Statistics in medicine 25, 3201–3216 (2006).

12 Haykin, S. & Network, N. A comprehensive foundation. Neural Networks 2 (2004).

13 Srivastava, N., Hinton, G. E., Krizhevsky, A., Sutskever, I. & Salakhutdinov, R. Dropout: a simple way to prevent neural networks from overfitting. Journal of Machine Learning Research 15, 1929–1958 (2014).

14 Harrell, F. E., Lee, K. L. & Mark, D. B. Tutorial in biostatistics multivariable prognostic models: issues in developing models, evaluating assumptions and adequacy, and measuring and reducing errors. Statistics in medicine 15, 361–387 (1996).

15 Huang, S., Yee, C., Ching, T., Yu, H. & Garmire, L. X. A Novel Model to Combine Clinical and Pathway-Based Transcriptomic Information for the Prognosis Prediction of Breast Cancer. PLoS computational biology 10, e1003851 (2014).

16 Huang, S. et al. Novel personalized pathway-based metabolomics models reveal key metabolic pathways for breast cancer diagnosis. Genome medicine 8, 1 (2016).

17 Gevrey, M., Dimopoulos, I. & Lek, S. Review and comparison of methods to study the contribution of variables in artificial neural network models. Ecological modelling 160, 249–264 (2003).

18 Olden, J. D., Joy, M. K. & Death, R. G. An accurate comparison of methods for quantifying variable importance in artificial neural networks using simulated data. Ecological Modelling 178, 389–397 (2004).

19 Broad. Broad Institute TCGA Genome Data Analysis Center (2014): Analysis Overview for 15 July 2014. Broad Institute of MIT and Harvard, doi:10.7908/C1DN43V9 (2014).

20 Love, M., Anders, S. & Huber, W. Differential analysis of RNA-Seq data at the gene level using the DESeq2 package. (2013).

21 Qian, N. On the momentum term in gradient descent learning algorithms. Neural networks 12, 145–151 (1999).

22 Bengio, Y., Boulanger-Lewandowski, N. & Pascanu, R. in Acoustics, Speech and Signal Processing (ICASSP), 2013 IEEE International Conference on. 8624–8628 (IEEE).

23 Battiti, R. Accelerated backpropagation learning: Two optimization methods. Complex systems 3, 331–342 (1989).

24 Wei, R. et al. Meta-dimensional data integration identifies critical pathways for susceptibility, tumorigenesis and progression of endometrial cancer. Oncotarget (2016).

25 Maaten, L. v. d. & Hinton, G. Visualizing data using t-SNE. Journal of Machine Learning Research 9, 2579–2605 (2008).

26 Subramanian, A. et al. Gene set enrichment analysis: a knowledge-based approach for interpreting genome-wide expression profiles. Proceedings of the National Academy of Sciences 102, 15545–15550 (2005).

27 Kanehisa, M. & Goto, S. KEGG: kyoto encyclopedia of genes and genomes. Nucleic acids research 28, 27–30 (2000).

28 Girgin, C. et al. P53 mutations and other prognostic factors of renal cell carcinoma. Urologia internationalis 66, 78–83 (2001).

29 Marques, I. et al. Influence of survivin (BIRC5) and caspase-9 (CASP9) functional polymorphisms in renal cell carcinoma development: a study in a southern European population. Molecular biology reports 40, 4819–4826 (2013).

30 Akhurst, R. J. & Derynck, R. TGF-β signaling in cancer—a double-edged sword. Trends in cell biology 11, S44–S51 (2001).

31 Choueiri, T. K. et al. Phase II and biomarker study of the dual MET/VEGFR2 inhibitor foretinib in patients with papillary renal cell carcinoma. Journal of Clinical Oncology 31, 181–186 (2013).

32 Cork, S. M. & Van Meir, E. G. Emerging roles for the BAI1 protein family in the regulation of phagocytosis, synaptogenesis, neurovasculature, and tumor development. Journal of molecular medicine 89, 743–752 (2011).

33 Fukushima, Y. et al. Brain-specific angiogenesis inhibitor 1 expression is inversely correlated with vascularity and distant metastasis of colorectal cancer. International journal of oncology 13, 967–970 (1998).

34 Lee, J. et al. Comparative study of angiostatic and anti-invasive gene expressions as prognostic factors in gastric cancer. International journal of oncology 18, 355–362 (2001).

35 Izutsu, T., Konda, R., Sugimura, J., Iwasaki, K. & Fujioka, T. Brain-specific angiogenesis inhibitor 1 is a putative factor for inhibition of neovascular formation in renal cell carcinoma. The Journal of urology 185, 2353–2358 (2011).

36 Nishimori, H. et al. A novel brain-specific p53-target gene, BAI1, containing thrombospondin type 1 repeats inhibits experimental angiogenesis. Oncogene 15, 2145–2150 (1997).

37 Kudo, S. et al. Inhibition of tumor growth through suppression of angiogenesis by brain-specific angiogenesis inhibitor 1 gene transfer in murine renal cell carcinoma. Oncology reports 18, 785–792 (2007).

38 Oka, H. et al. Constitutive activation of mitogen-activated protein (MAP) kinases in human renal cell carcinoma. Cancer research 55, 4182–4187 (1995).

39 Friday, B. B. & Adjei, A. A. Advances in targeting the Ras/Raf/MEK/Erk mitogen-activated protein kinase cascade with MEK inhibitors for cancer therapy. Clinical Cancer Research 14, 342–346 (2008).

40 Nguyen, T.T., Huang, J. Z. & Nguyen, T. T. Unbiased Feature Selection in Learning Random Forests for High-Dimensional Data. The Scientific World Journal 2015 (2015).

41 LeCun, Y. & Bengio, Y. Convolutional networks for images, speech, and time series. The handbook of brain theory and neural networks 3361, 1995 (1995).

